# Differential impacts of repeated sampling on odor representations by genetically-defined mitral and tufted cell subpopulations in the mouse olfactory bulb

**DOI:** 10.1101/2020.03.03.975151

**Authors:** Thomas P. Eiting, Matt Wachowiak

## Abstract

Sniffing—the active control of breathing beyond passive respiration—is used by mammals to modulate olfactory sampling. Sniffing allows animals to make odor-guided decisions within ~200 ms, but animals routinely engage in bouts of high-frequency sniffing spanning several seconds; the impact of such repeated odorant sampling on odor representations remains unclear. We investigated this question in the mouse olfactory bulb, where mitral and tufted cells (MTCs) form parallel output streams of odor information processing. To test the impact of repeated odorant sampling on MTC responses, we used two-photon imaging in anesthetized male and female mice to record activation of MTCs while precisely varying inhalation frequency. A combination of genetic targeting and viral expression of GCaMP6 reporters allowed us to access mitral (MC) and superficial tufted cell (sTC) subpopulations separately. We found that repeated odorant sampling differentially affected responses in MCs and sTCs, with MCs showing more diversity than sTCs over the same time period. Impacts of repeated sampling among MCs included both increases and decreases in excitation, as well as changes in response polarity. Response patterns across ensembles of simultaneously-imaged MCs reformatted over time, with representations of different odorants becoming more distinct. MCs also responded differentially to changes in inhalation frequency, whereas sTC responses were more uniform over time and across frequency. Our results support the idea that MCs and TCs comprise functionally distinct pathways for odor information processing, and suggest that the reformatting of MC odor representations by high-frequency sniffing may serve to enhance the discrimination of similar odors.

## Introduction

Olfaction, a key sensory modality for mammals, has emerged as a leading system in which to investigate how active sensation contributes to sensory processing (Uchida et al., 2006; Wachowiak, 2011; Jordan et al., 2018b). Sniffing—the active control of breathing beyond passive respiration—is used by mammals to modulate olfactory sampling. Sniffing provides temporal control over olfactory sensation and can increase access to odor information by increasing the flow of odorants through the nasal cavity, and also by allowing multiple samples of odorant in a shorter time (Zhao et al., 2006; Verhagen et al., 2007; Courtiol et al., 2011; Rygg et al., 2017). Repeated high-frequency (3 – 10 Hz) inhalations are a hallmark of sniffing behavior in most mammals, including rodents (Youngentob et al., 1987; Laska, 1990; Thesen et al., 1993; Wesson et al., 2008a). While mice and rats can discriminate odors in as little as one sniff (150-200 ms (Uchida and Mainen, 2003; Abraham et al., 2004), they frequently exhibit longer sniffing bouts lasting several seconds (Welker, 1964; Wesson et al., 2008a). These longer-term inhalation dynamics may play a role in more complex olfactory tasks, such as matching an odor to a known odor object, gleaning precise concentration information, or evaluating complex mixtures.

How repeated sampling influences olfactory processing by neural circuits in the olfactory bulb (OB) or elsewhere is not well understood. Previous work has shown that rapid sniffing induces adaptation of olfactory sensory inputs to the OB (Verhagen et al., 2007); more recently, we and others have found that inhalation frequency modulates subthreshold membrane potential and spiking patterns in mitral and tufted cells (MTCs; (Bathellier et al., 2008; Carey and Wachowiak, 2011; Díaz-Quesada et al., 2018; Jordan et al., 2018b)). Multiple distinct functions of this frequency-dependent modulation have been proposed, including enhancing olfactory sensitivity, increasing the salience of odorants against a background, and improving fine-scale odor discrimination.

There is also substantial evidence that mitral cells (MCs) and tufted cells (TCs) have distinct odorant response properties and likely contribute differentially to encoding odor information (Nagayama et al., 2004; Griff et al., 2008; Fukunaga et al., 2012; Jordan et al., 2018a; Short and Wachowiak, 2019). For example, some studies have reported that MCs appear more subject to inhibition, respond with longer latency relative to inhalation, and may hyperpolarize more frequently than TCs during bouts of high-frequency inhalation (Jordan et al., 2018b). Such differences have led to the suggestion that TCs convey rapid odor “snapshots,” whereas MCs contribute more complex, time-varying information (Abraham et al., 2004; Abraham et al., 2010; Najac et al., 2011; Fukunaga et al., 2012).

Ultimately, because odor coding involves combinatorial activity patterns across many MTCs, to understand how active sniffing impacts odor representations among MCs and TCs it is important to examine its impacts on populations of multiple MTCs recorded simultaneously. With a few exceptions (e.g., Patterson, Lagier, & Carleton, 2013), prior studies have relied on recordings from one or a small number of simultaneously-recorded cells. Here, we investigated the contributions of inhalation frequency in shaping odor representations by MCs and TCs in anesthetized mice, using two-photon imaging from genetically- and anatomically-defined subsets of MCs and TCs, imaging from multiple cells simultaneously. We found notable population-level differences between MCs and a subset of TCs defined by their superficial location and their expression of the peptide transmitter cholecystokinin (CCK). The CCK+ superficial TC (sTC) population showed stronger coupling to individual inhalations, few suppressive responses, and little change in its population response pattern during repeated odorant sampling. In contrast MCs showed diverse changes in excitation and suppression with repeated odorant sampling, such that the MC population response pattern reformatted substantially from the beginning to the end of the odorant presentation, and did so in a frequency-dependent manner. These effects point to an important role for active sensing in odor perception and highlight the importance of considering neural activity dynamics over a time-scale beyond that of a single inhalation.

## Materials and Methods

### Animals

Adult transgenic mice (ages 6-16 weeks) of both sexes were used in all experiments. Imaging in mice was carried out in Cck-IRES-cre (Jackson Laboratory (Jax) stock #012706) (Taniguchi et al., 2011) mice for imaging in sTCs or in either Pcdh21-cre (Mutant Mouse Resource and Research Center stock #030952-UCD) (Gong et al., 2003) mice or Tbet-cre (Jax stock #024507) (Haddad et al., 2013) mice for imaging in MCs. Mice were either crossed with Ai95(RCL-GCaMP6f)-D (Jax stock #024105) (Madisen et al., 2015) reporter mice or injected with adeno-associated viral vectors (AAV) to drive expression (see below). All procedures were carried out according to NIH guidelines and were approved by the Institutional Animal Care and Use Committee of the University of Utah.

### Viral Vector Expression

GCaMP6f/s expression was achieved using recombinant viral vectors, AAV1, AAV5, or AAV9 serotypes of hSyn.Flex.GCaMP6f.WPRE.SV40 or hSyn.Flex.GCaMP6s.WPRE.SV40 (UPenn Vector Core), injected using pulled glass pipettes into the anterior piriform cortex (aPC) at the following coordinates (relative to Bregma): 2.4 mm anterior, 1.6 mm lateral, and 3.5 mm ventral. Injections were performed as described previously (Rothermel et al., 2013; Wachowiak et al., 2013). Animals were injected with carprofen (5 mg/kg; analgesic) and enrofloxacin (10 mg/kg; antibiotic) immediately prior to surgery as well as the following day. After surgery, animals were singly-housed and were allowed to recover on a heating pad prior to transfer back to the animal colony. Imaging experiments occurred 14-45 days after injection.

### Olfactometry

Odorants were delivered via an air-dilution olfactometer, in which the odorants were first diluted in mineral oil, followed by an air-phase dilution of nitrogen-diluted odorant into normal air, for a final estimated concentration presented to the animal of ~10-20 ppm (Wachowiak et al., 2013; Economo et al., 2016). Odorants of different chemical groups (esters, aldehydes, ketones, organic acids) were presented in random order with respect to inhalation frequency, and repeated 8 times with a minimum of 36 second inter-trial interval (ITI; Fig. 1). Inhalation (1, 3, and 5 Hz) was controlled using an artificial inhalation paradigm (Wachowiak and Cohen, 2001; Díaz-Quesada et al., 2018; Eiting and Wachowiak, 2018). Stimuli and artificial inhalation were controlled with Labview software (National Instruments).

**Figure 1.**
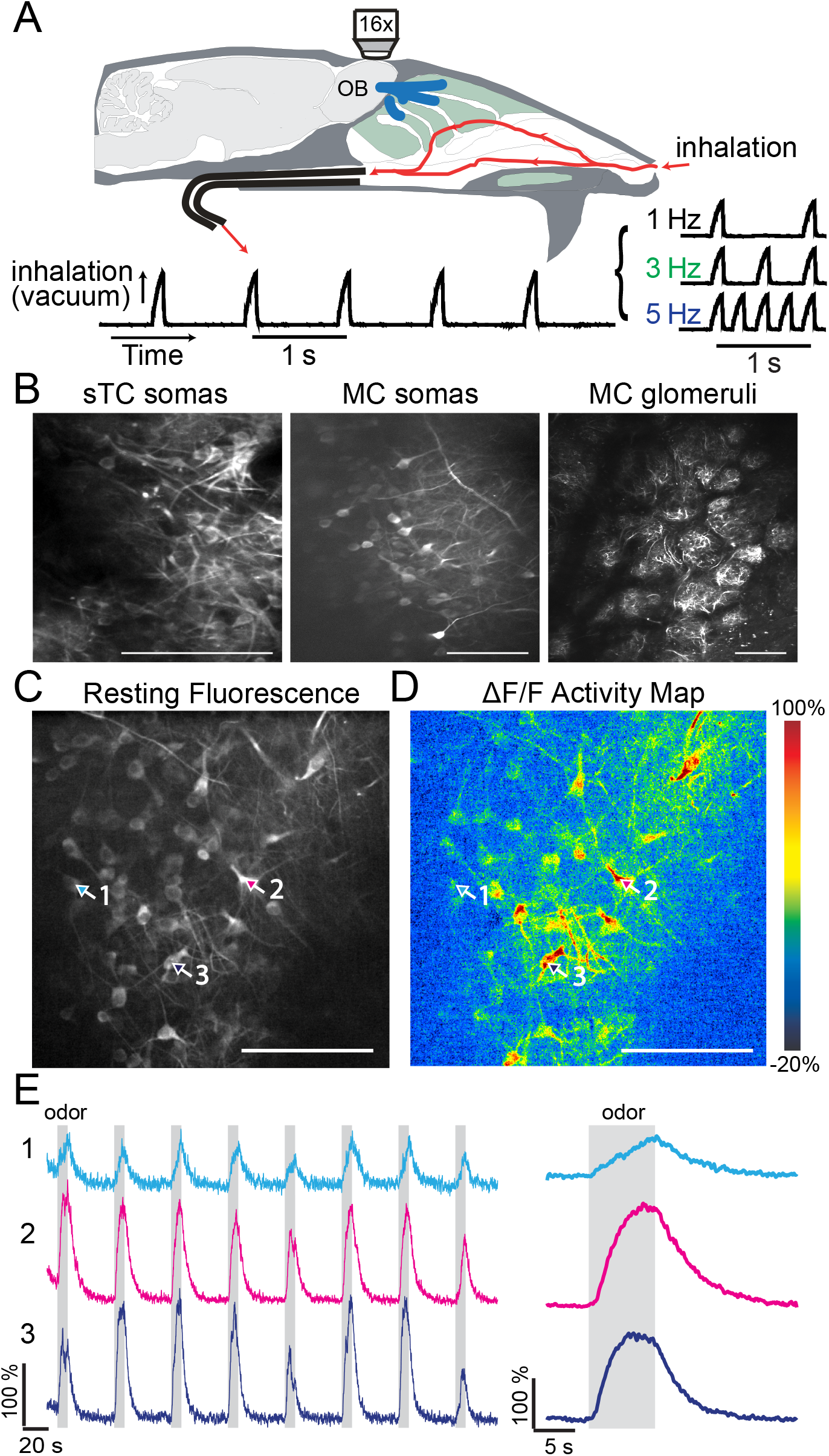
Imaging mitral and tufted cell population odor responses with artificial inhalation. ***A***, Schematic of experimental paradigm. Odorants are delivered using an artificial inhalation paradigm, with inhalation frequencies of 1, 3, or 5 Hz. A sample inhalation trace (1 Hz) is pictured at the bottom. ***B,*** Representative images of resting fluorescence in three types of experiments, showing typical fields of view for sTC somas, MC somas, and MC-innervated glomeruli. ***C*,** Resting fluorescence image from one field of view, showing over 50 MC somas. ***D*,** Odorant-evoked calcium signals, represented as ΔF/F, of the same field of view shown in ***C***. ***E*,** Left, Example traces (from ROIs indicated in ***C*** and ***D***), showing evoked fluorescence for 8 consecutive trials in which odorant is presented during 5 Hz inhalation for 8 s per trial. Right, average responses of the three ROIs. For most analyses, average responses were used in calculations. All scale bars = 100 μm.

### In vivo two photon imaging

Animals were initially anesthetized with pentobarbital (50 mg/kg); and long-term anesthesia was maintained with isoflurane (0.5%-1.0% in oxygen) for the duration of the experiment. Body temperature was maintained at 37°C. Following a craniotomy (~1.5 x 1 mm) over one olfactory bulb, we imaged glomeruli (MCs) or cell bodies (sTCs and MCs) at appropriate depth: 20-30 μm below olfactory nerve layer (ONL) for glomeruli, ~120-150 μm below the ONL for sTCs and ~240-270 μm below ONL for MCs. Calcium signals (GCaMP6f/6s) were collected using a 2-photon laser and microscope system (Neurolabware), running a Ti-Sapphire laser (Coherent, Chameleon Ultra-II) operating at 940 nm. Imaging was performed through a 16x, 0.8 NA objective (Olympus) and emitted light was collected through a GaAsP photomultiplier tube (Hamamatsu, H10770B-40). Frame rate was 15.5 Hz.

### Data Analysis

Image processing and analysis of optical signals followed procedures similar to those described previously (Wachowiak et al., 2013; Economo et al., 2016). Analyses of calcium data were performed on fluorescence signals collected at 15.5 Hz. Regions of interest (ROIs) were either glomeruli (MCs) or cell bodies (MCs and sTCs), selected as contiguous polygons of responsive pixels. ROIs were initially defined based on visually inspecting average odor-evoked response maps and choosing a single polygon above a certain visual threshold of response relative to background. Some ROIs were selected using a manual procedure to select similarly strong responses. ΔF/F values were calculated as the change in fluorescence during 8 s odor stimulation compared to the 4 s immediately prior to odor stimulation, divided by the mean baseline fluorescence value. ROIs were included for final analyses based on a criterion of the peak value of the ΔF/F signal reaching above 6 standard deviations of the signal during the 4 s immediately prior to the odorant stimulation. Analyses were carried out only when blank air generated few or no responses (max 5% responding cells/glomeruli), and individual ROIs were removed from analyses if they responded to blank air stimulation. Analyses of the population correlation at a given time point to the activity within the first time point (Figs. 3*C*,*D*, Fig. 5*C*) were carried out on glomerular or soma population activity across each odor-frequency combination per animal, then grouped together either by experiment or by cell type, as indicated in the Results.

**Figure 2.**
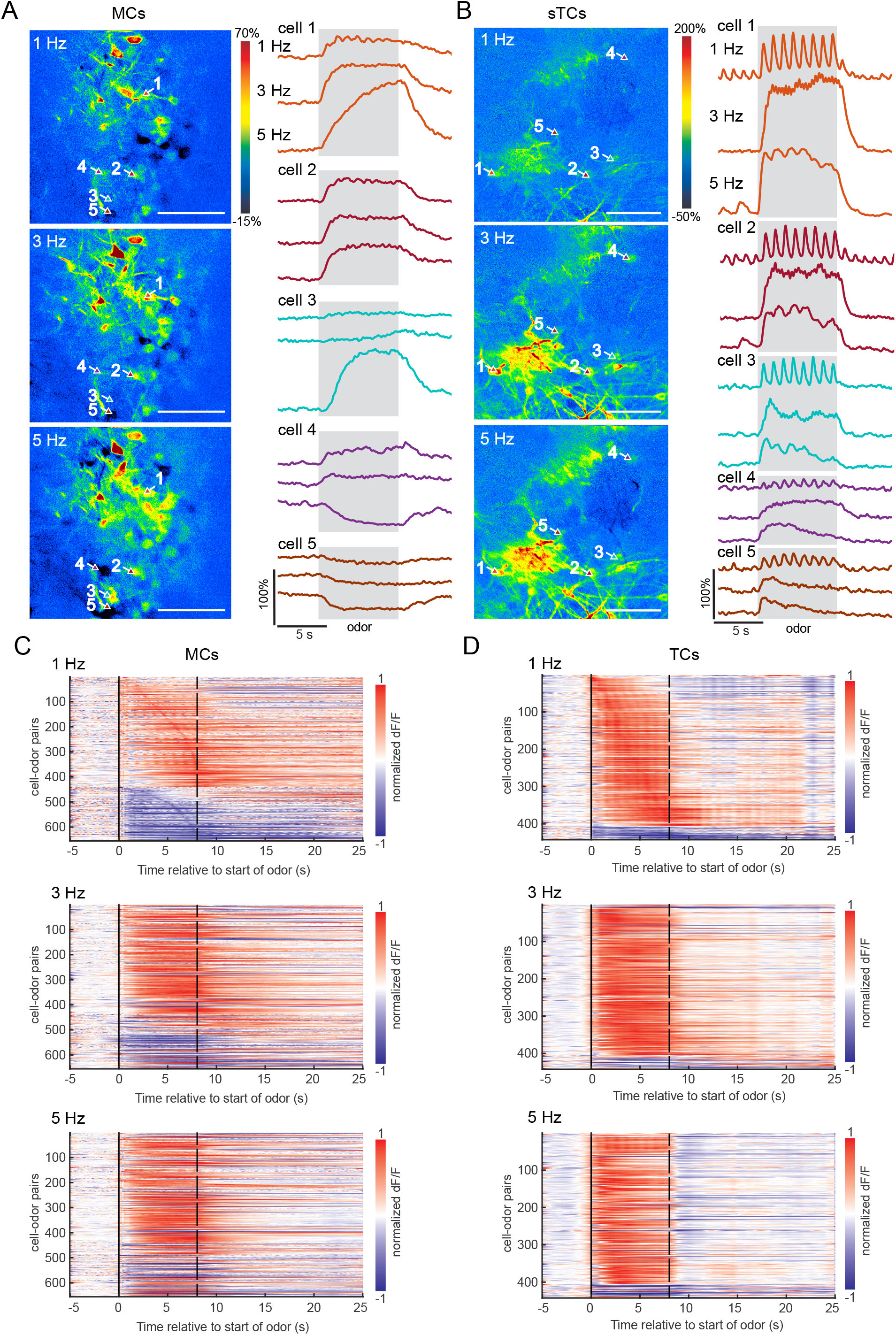
Odorant-evoked activity varies with inhalation frequency. ***A***, Example trialaveraged responses from five MC somas, showing diverse responses to ethyl butyrate (0.1%) at three different inhalation frequencies. ***B*,** Trial-averaged responses from five sTC somas, showing a relative lack of diversity in temporal responses among cells within a given preparation. In all cells, fluorescence signals clearly follow 1 Hz inhalations (top rows for each cell), but this effect disappears at higher frequencies. Scale bars for the maps in ***A*** and ***B*** = 100 μm. ***C,*** Time-by-activity plots for all MC cell-odor pairs, rank-ordered by time-to-peak activity at 1 Hz. MC responses are diverse and prolonged, and activity patterns change substantially among cells at higher inhalation frequency. ***D*,** Time-by-activity plot for sTC somas. Note that, compared to ***C***, peak sTC activity tends to occur earlier than peak MC activity, and sTC responses generally turned off quickly after odor presentation compared to MC responses. In both ***C*** and ***D***, odorant offset is at 8 sec.

**Figure 3.**
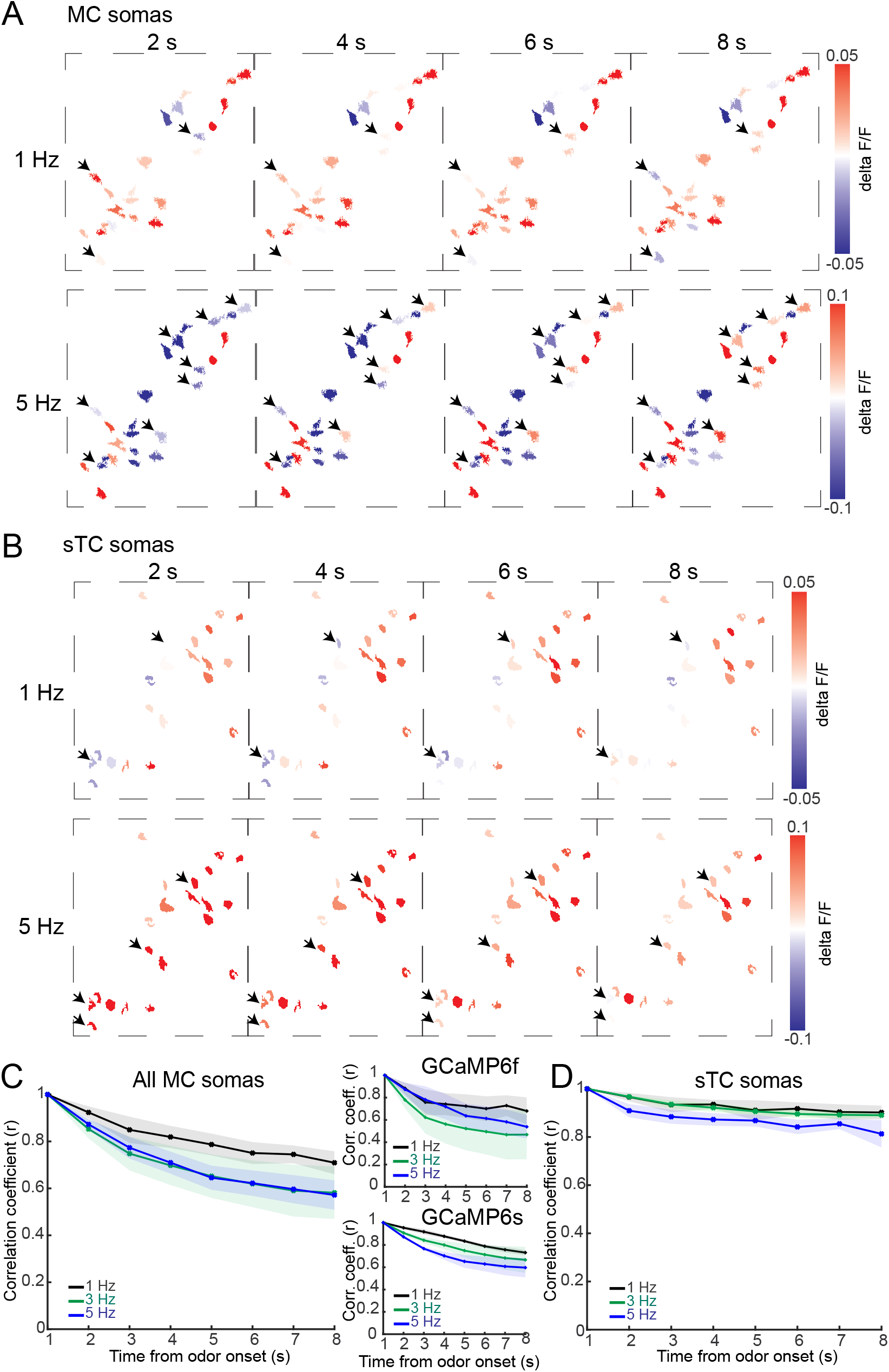
MC population responses evolve with repeated odorant sampling more than sTC populations. ***A**,* Example MC soma experiment, showing responsive cells during1 Hz and 5 Hz inhalation at different time-points. Time progresses horizontally in 2 s increments that represent the response maps during odorant presentation, averaged over 1 s. A number of MC somas change their response strength or response polarity during 5 Hz inhalation, with fewer changing during 1 Hz inhalation. Black arrows highlight cells whose response changes substantially or switches polarity during odor presentation. ***B,*** Example sTC soma experiment, displayed as in (*A*), showing a relative lack of changes in responses among individual cells, regardless of inhalation frequency. ***C,*** Average MC population response to an odorant at a given time point correlated to the population response at the first time point over the length of odorant presentation. Correlations decreased faster at higher frequencies, though this difference was not statistically significant. Left, all MC cell body data. Right top, data from GCaMP6f experiments; right bottom, data from GCaMP6s experiments. ***D*,** Plot of same analysis for sTC responses. Temporal decorrelation does not vary substantially across inhalation frequencies among sTCs. Decorrelation values differed between MCs and sTCs (see text for details). In both ***C*** and ***D***, shaded areas represent SEM.

**Figure 4.**
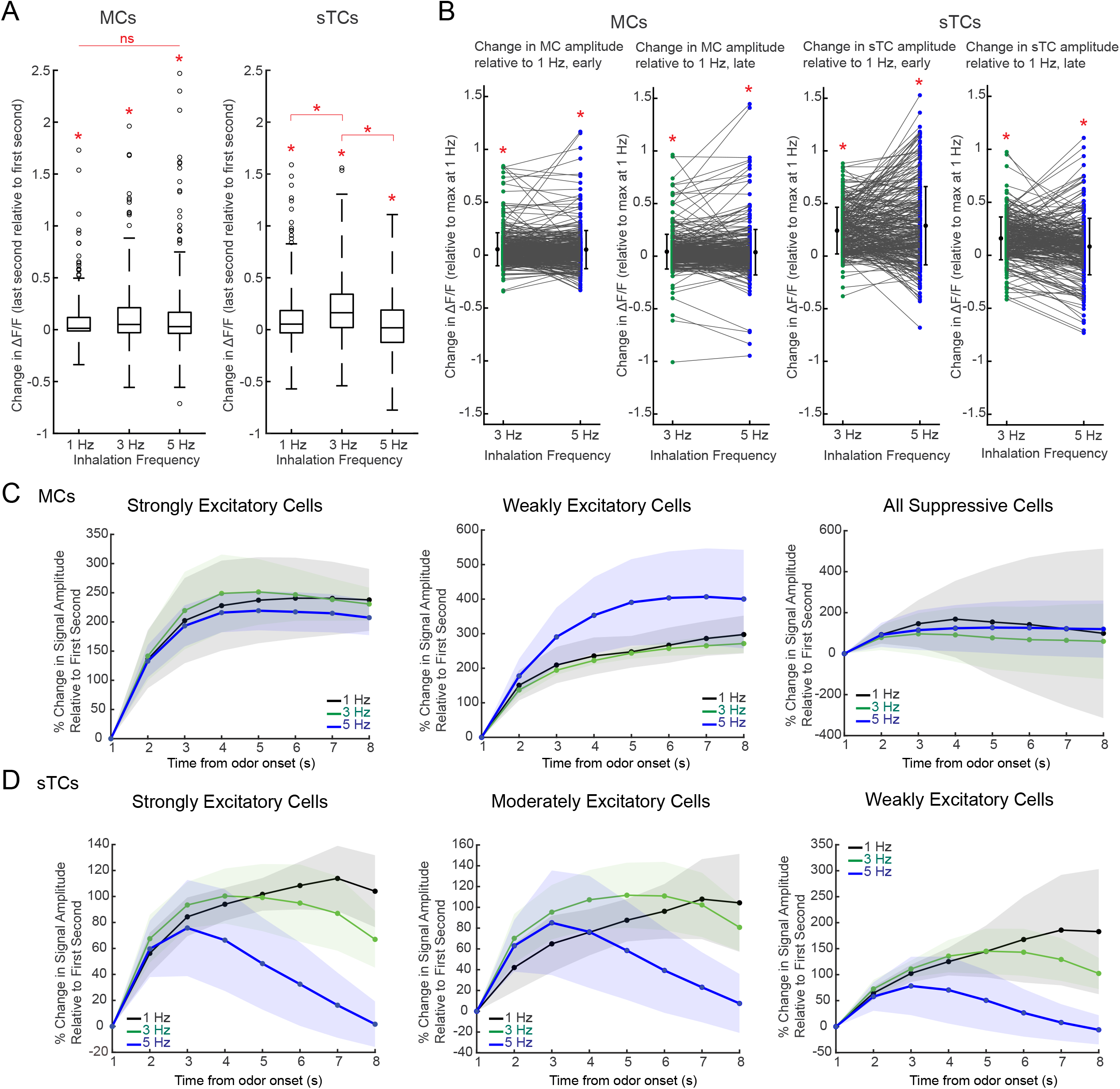
MC and TC responses show high variance in response to changing inhalation frequency. ***A**,* Change in signal amplitudes (ΔF/F) from the last second to the first second for all three inhalation frequencies across both cell types. In both MCs and sTCs, responses late in the odor presentation (final 1 s) are slightly greater than responses early in the odor presentation (first 1 s), but mean changes are small relative to variance across cells. Boxplots represent means and 25%-75% percentiles, and whiskers extend for 3 SDs. Outliers are plotted as individual points outside of the whiskers. ***B*,** Change in strength of response magnitudes at 3 Hz and 5 Hz compared to 1 Hz, for both MCs (left) and sTCs (right). For both cell types, the plot on the left reflects change within the first second of odorant presentation, and the plot on the left shows change within the last second of odorant presentation. Points in each plot are relative to maximum value at 1 Hz. For both MCs and sTCs, higher-frequency inhalations lead to increased excitation (means in each plot are greater than zero). For MCs, early amplitudes change less than late amplitudes. The same holds for sTCs, though early amplitude changes are relatively much higher compared to MCs. Activity within the same cells are joined by lines. Beside the individual points, mean ± SEM is plotted for each case. ***C*,** Change in response amplitude over time for three groups of MCs, plotted as % change relative to the amplitude within the first 1 s time bin. Changes in response amplitudes of strongly excitatory and suppressed cells show little divergence between high and low frequency inhalation, whereas weakly excitatory cells show a notable increase in excitation at 5 Hz inhalation. ***D*,** Same analysis as in ***C***, plotted for strongly, moderately and weakly-excited sTCs. Here, all groups respond in a similar fashion to inhalation frequency. In general, responses peak earlier at high inhalation frequencies compared to lower frequencies, and the relative pattern within a grouping of cells stays consistent across groups. In both ***C*** and ***D***, shaded areas represent SEM. *= *p* < 0.01 (see text for specific values).

**Figure 5.**
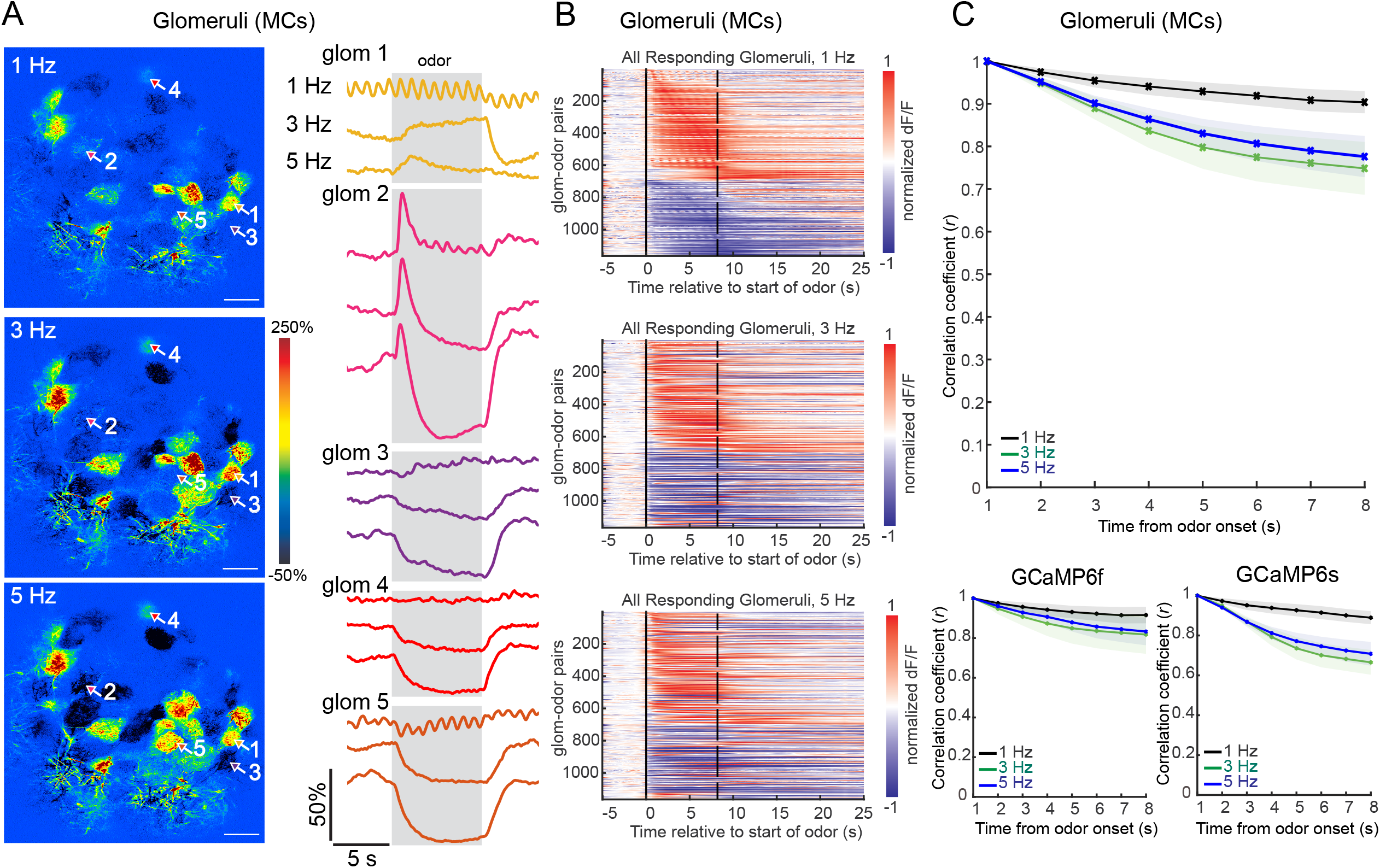
Glomerular signals from apical dendrites of MCs show similar diversity in responses and decorrelation patterns compared to MC cell somas. ***A***, Field of view of one experimental preparation, showing how the same set of glomeruli respond to three different inhalation frequencies of the same odorant. Right, example trial-averaged responses from five glomeruli. Glomeruli have variable responses at differing inhalation frequencies. Glomerulus 2, for example, shows a quick burst of activity at all three frequencies, but at 3 and 5 Hz, this initial burst is followed by a rapid decrease in activity below baseline, which is not seen at 1 Hz. Glomerulus 4 shows slight excitation at 1 Hz, and gradual suppression at 3 and 5 Hz. Scale bars = 100 μm. ***B,*** Time-by-activity plots for all glomeruli, rank-ordered by time-to-peak activity at 1 Hz. Similar to MC somas, glomerular responses are diverse and prolonged. MC glomeruli include a high proportion of suppressive responses. ***C*,** Correlation of the average glomerular population response to an odorant at a given time point to the population response at the first time point, over the length of odorant presentation. Top, all MC glomerular data. Bottom left, data from GCaMP6f experiments; bottom right, data from GCaMP6s experiments. MC glomerular responses decorrelate more strongly at 3 and 5 Hz compared to 1 Hz. The magnitude of decorrelation is lower than in MC soma data (compare to Fig. 3A).

Principal components analysis (PCA) was carried out for experiments in which three or more odorants generated consistent responses, resulting in 5 MC-soma experiments and 3 sTC-soma experiments (Fig. 6). For each experiment, PCA was carried out as follows. First, responses to any given odor-by-inhalation frequency condition were averaged across the eight repeated presentations of that stimulus, and binned per frame (i.e. ~15.5 Hz; each bin ~65 s). Data were then concatenated for all odors into one dataset for each inhalation-frequency condition. PCA was carried out on each of these three datasets (1, 3, and 5 Hz), using only the first three PCs in each case. Euclidean distances were calculated between each successive bin and summed across all odors for each inhalation condition. Euclidean distances were standardized by number of odorants within a given experiment, because this measure provides an unbiased method to compare the spread of points in PC space across experiments or conditions.

**Figure 6.**
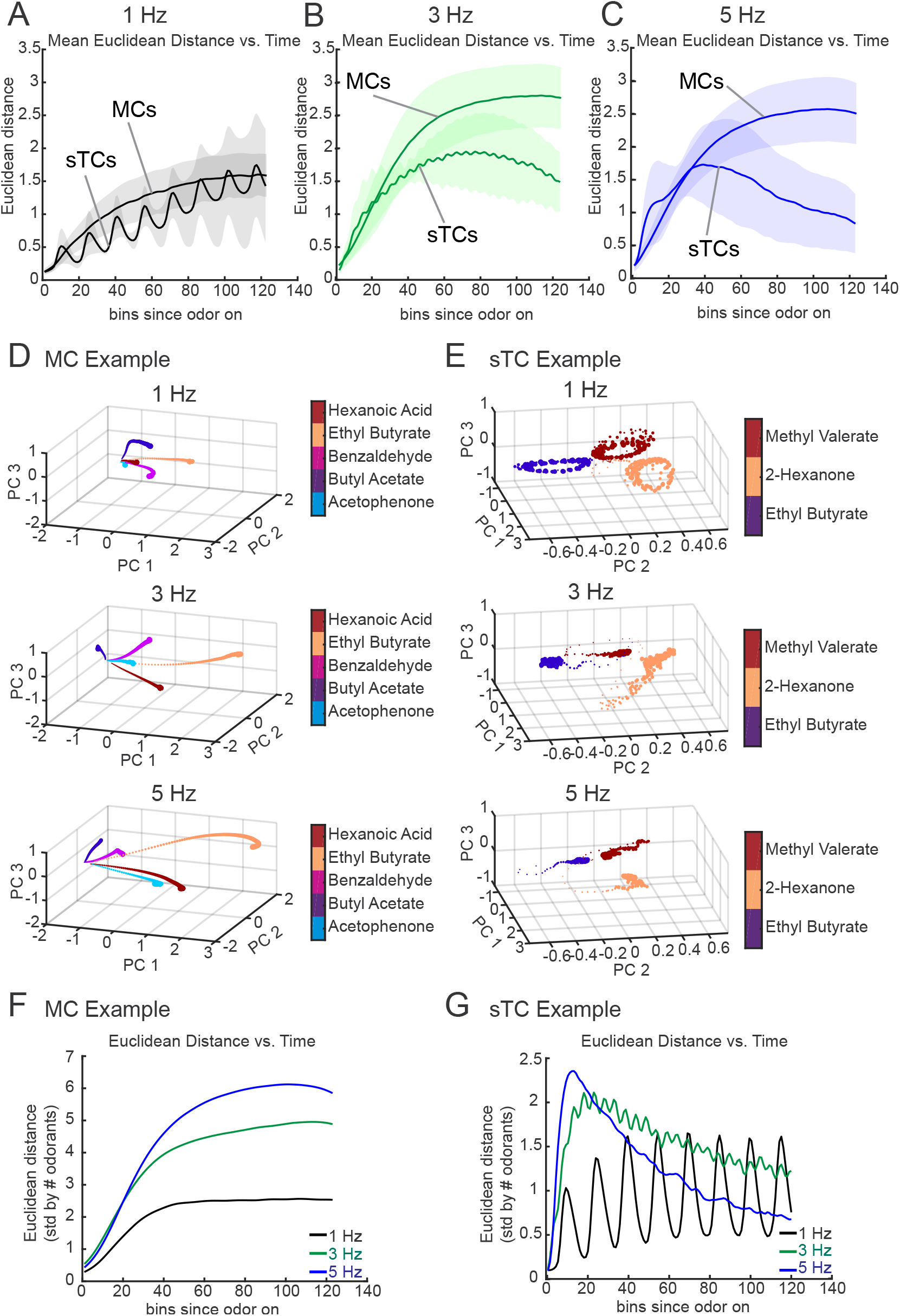
Odor representations become more distinct over time at high inhalation frequencies in MCs but not in sTCs. ***A**,* Summed Euclidean distances across MC and sTC preparations at 1 Hz inhalation frequency, showing that the distances parallel each other in both cell types. *B*, Euclidean distance at 3 Hz exhibit different trends between MCs and TCs, diverging midway through the 8 s odor presentation. *C,* At 5 Hz inhalation, distances diverge between MCs and sTCs after roughly 3 seconds of odorant presentation. In *A-C*, shaded areas represent SEM. *D,* Euclidean distance over time for MCs, showing that odorant representations at higher frequencies become more separated within the same 8 s odorant presentation period as the 1 Hz condition (1, 3 and 5 Hz data shown from top to bottom). Thus, each odorant becomes more unique in terms of the MC population response that it generates. *E,* Euclidean distance over time for sTCs. These plots show little differentiation among the three inhalation frequencies. Instead, the dominant feature visible throughout all inhalation frequencies is of cyclic fluctuations in odorant population representations corresponding to each inhalation. *F,* Example MC experiment, showing a rapid increase in distances at the beginning of odorant presentation (especially at 3 and 5 Hz), followed by a slow rise/plateauing of Euclidean distance for the bulk of the odorant presentation. *G*, Example sTC experiment, showing that one of the dominant longer-term signals is that sTC responses closely follow inhalation, at least at low frequencies. Odorant trajectories do not spread out as much at high frequencies as compared to the MC case, and a major feature is the oscillatory nature of the PCA, reflecting the inhalation-linked responses of sTCs.

Analyses and statistical tests were carried out using custom scripts written in Matlab 2019b (Mathworks) or *R* v.3.3.2 (R Foundation for Statistical Computing). Raw data and analysis code are available upon request.

## Results

Our major goals were to understand how inhalation frequency affects activity patterns of mitral and tufted cell populations, and to explore differences in their responses in the context of inhalation frequency. To achieve these aims we collected fluorescent calcium signals from mitral (MC) and superficial tufted (sTC) cells under 2-photon microscopy using an artificial-inhalation paradigm to control inhalation frequency in isoflurane-anesthetized mice (Fig. 1*A, B*)(Díaz-Quesada et al., 2018; Eiting and Wachowiak, 2018). To target MCs, protocadherin21-Cre mice were injected with cre-dependent virus into the anterior piriform cortex to generate expression predominately in prifiorm-projecting MCs (Rothermel et al., 2013). To target superficial tufted cells (sTCs), we crossed cholecystokinin (CCK)-IRES-Cre mice with Rosa-GCaMP6f mice (line Ai95D), as previously reported (Economo et al., 2016; Short and Wachowiak, 2019). For each cell type, we lowered the imaging plane to the appropriate depth. We gathered MC and sTC responses from populations of cells imaged while presenting clean air and then odorant at different inhalation frequencies (see Methods). We presented eight successive odorant presentations of 8 s duration (interspersed with a 36 s inter-trial interval) at each inhalation frequency. Example resting fluorescence, activity map, and traces from a MC-soma imaging experiment show examples of average traces produced from individual trials (Fig. 1C-*E*).

We first compared responses of MCs and sTCs to repeated sampling of odorant at 1, 3, and 5 Hz. Examples of imaging experiments targeted to each cell type are shown in Figs. 2*A* and *B*, illustrating qualitative differences in how MCs and sTCs respond to repeated sampling, and how these responses change as a function of frequency. MCs showed more diverse temporal responses over time that included slower increases in activity peaking over the course of odorant presentation, while sTC responses tended to be simpler, generally following the onset and offset of odorant presentation (Fig. 2*A*, *B*). sTCs also more commonly followed individual inhalations at 1 Hz than did MCs. These trends were evident across the entire population of imaged MCs and sTCs, compiled from seven and five mice, respectively (Fig. 2*C*, *D*). Finally, suppressive responses were common in MCs but were rarely observed in sTCs. MC responses included a substantial percentage of suppressive cells: 123/450 (27%) cell-odor pairs at 1 Hz, 135/580 (23%) cell-odor pairs at 3 Hz, and 147/612 (24%) cell-odor pairs at 5 Hz were suppressive. By contrast there were very few suppressive sTC responses: 8/233 (3-4%) cell-odor pairs at 1 Hz, 8/265 (3%) cell-odors pairs at 3 Hz, and 4/269 (1-2%) cell-odor pairs at 5 Hz were suppressive. Signals from MC somas using both GCaMP6f (4 mice) and GCaMP6s (3 mice) showed similar temporal response patterns and prevalence of suppression (not shown), suggesting that the observed differences between MC and sTC populations were likely not due to differences in the temporal dynamics of the calcium indicator used. Overall, this suite of differences recapitulates those reported earlier at 2 Hz inhalation frequency (Economo et al., 2016).

MCs could respond heterogeneously to different inhalation frequencies (Fig. 2*A*, *B*) – for example, cell 3 in Fig. 2*A* shows no response at 1 Hz inhalation but a strong excitation at 5 Hz, while cell 4 switches response polarity from excitation at 1 and 3 Hz to suppression at 5 Hz. Across the MC population, cells switched response polarity from suppressed to excited roughly 3-4 times as frequently as excited to suppressed: 49/438 (11%) excited to suppressed from 1 Hz to 3 Hz, 64/438 (15%) excited to suppressed from 1 Hz to 5 Hz; 84/217 (39%) suppressed to excited from 1 Hz to 3 Hz, 111/217 (51%) suppressed to excited 1 Hz to 5 Hz (Fig. 2*C*). By contrast, sTCs typically responded similarly to all three inhalation frequencies (Fig. 2*B*, *D*), although in many cells a decrease in excitation suggesting adaptation was apparent with repeated sampling at higher frequencies (e.g., cells 4 and 5 in Fig. 2*B*). Changes in response polarity in sTCs were much more rare compared to MCs: 22/405 (5%) switched from excited to suppressed responses when comparing 1 Hz to 3 Hz, and 17/405 (4%) switched from excited to suppressed in the 1 Hz to 5 Hz case. Furthermore, while only 3-4% of sTCs showed suppressive responses to 1 Hz inhalation, nearly half – 41% (15/37) and 46% (17/37) – of these cells switched polarity from suppression to excitation when inhalation frequency increased to 3 Hz and 5 Hz, respectively. Overall, these analyses indicate that MCs and sTCs show distinct impacts of inhalation frequency on responses to repeated odorant inhalation, but that across both cell types, as inhalation frequency increased, cells tended to switch from suppression to excitation much more frequently than from excitation to suppression.

Heterogeneity in MC (and to a lesser extent sTC) responses implies that neural representations of odor identity, measured in terms of spike rate across each population, will change as a function of repeated odor sampling (Patterson et al., 2013; Díaz-Quesada et al., 2018). Example experimental preparations in Fig. 3*A*, *B* show how populations of cells (MCs in Fig. 3*A*, sTCs in Fig. 3*B*) change over an 8 s odorant presentation at 1 Hz and 5 Hz inhalation. In the MC example, numerous somas change their response polarity or their response amplitude across the 8 s odorant period, and this shift occurs much more frequently at higher inhalation frequency (Fig. 3*A*). By contrast, in the sTC example, very few cells switch their response polarity, or even change their activity level appreciably, regardless of inhalation frequency (Fig 3*B*).

To quantify changes among populations of MCs and sTCs, we correlated the population response to an odorant at a given time point to the population response at the first time point (corresponding to the first inhalation at 1 Hz, see Methods for precise times) over the length of odorant presentation. This analysis was performed separately for each ensemble of MCs or sTCs imaged in the same field of view (n = 13-58 MCs and 10-43 sTCs per field of view, 1 field of view per animal). Correlation series for different odorants tested in the same field of view were averaged before compiling across animals, to avoid bias from different numbers of odorants tested in different animals. These analyses revealed substantial differences in the evolution of odor representations between MCs and sTCs. Among MCs (Fig. 3*C*), high-frequency inhalation resulted in faster and more substantial decorrelation compared to low-frequency inhalation. After 8 s, the correlation to the initial response pattern was relatively high at 1 Hz (*r* = 0.71), and noticeably lower at higher inhalation frequencies (3 Hz, *r* = 0.58; 5 Hz, *r* = 0.57), though these differences were not statistically significant (n = 7 mice, Mann-Whitney U-test), in part because of the large heterogeneity in individual responses which led to high variance in decorrelations (see below). The qualitative result for MCs contrasts with that for sTCs (Fig. 3*D*), which maintained highly similar response patterns over the 8 s odor duration, with correlations to the initial response of *r =* 0.90 in the 1 Hz case, 0.89 at 3 Hz, and 0.81 Hz at 5 Hz (n = 5 mice). Regardless of inhalation frequency, on average, MCs decorrelated significantly more over the 8 seconds compared to sTCs (t-test, *p* = 0.0193). MC somas decorrelated to similar levels when either GCaMP6s or GCaMP6f was used (Fig. 3*C*), indicating that the differences observed between MC and sTC populations were not a result of different GCaMP reporters; in subsequent analyses we combined data from both variants. Overall, this analysis shows that MC responses show more decorrelation in response patterns with repeated sampling than do sTCs, and suggests that higher inhalation frequencies drive greater decorrelation in the MC population.

Linear decorrelation in population response patterns could arise from systematic changes in MC (or sTC) responsiveness. Such changes could include, for example, increases in excitation that saturate at the high end of a cell’s dynamic range or rise above threshold at the low end, or decreases in excitation that have the opposite effects. Alternatively, decorrelation could result from distinct effects of repeated sampling in different cells. To distinguish between these possibilities, we measured the change in MC or sTC excitation over the course of the odorant presentation, expressed as the change in ΔF/F from the first to the last second of the presentation (Fig. 4*A*). There was substantial heterogeneity in the effect of repeated sampling on excitation, reflected as a large variance across cells in the change in ΔF/F (Fig. 4*A*). Across all cells analyzed and all inhalation frequencies, there was a significant net increase in ΔF/F from the first to the last second, though the magnitude of the difference was small relative to the variance across different cells (Fig. 4*A*; Wilcoxon sign-ranked test: MC, 1 Hz, mean change in ΔF/F ± SD = 0.075 ± 0.22, *p* = 1.2×10^−11^; 3 Hz, 0.13 ± 0.31, *p* = 1.1×10^−15^; 5 Hz, 0.12 ± 0.36, *p* = 3.1×10^−10^; sTC, 1 Hz, 0.12 ± 0.31, *p* = 7.5×10^−11^; 3 Hz, 0.21 ± 0.30, *p* = 3.5×10^−26^; 5 Hz, 0.056 ± 0.29, *p* = 3.1×10^−2^). The magnitude of these differences did not vary significantly across inhalation frequencies among MCs, though it did among sTCs (Kruskal-Wallis test: MC, *χ^2^* = 5.12,1176, *p* = 0.08; sTC, *χ^2^* = 49.82,813, *p* = 1.5×10^−11^), with the sTC responses at 3 Hz increasing significantly more than at 1 Hz or 5 Hz inhalation frequencies (bonferroni-adjusted pairwise comparisons, *p* <0.00001 for both 1 Hz-vs-3 Hz and 3 Hz-vs-5 Hz).

We next asked whether increasing inhalation frequency systematically impacted MC or sTC excitability within a given cell, either initially (i.e., in the first second) or after prolonged sampling (i.e., after 8 seconds). To test this we compared response magnitude (ΔF/F) for the same cell-odor pairs at 1 Hz inhalation with that at 3 and 5 Hz, for both the initial (i.e., ‘early’) and 8-second (i.e, ‘late’) time-points (Fig. 4*B*). We found that increasing inhalation frequency significantly increased excitation for both early and late time-points in both MCs and sTCs (Fig.4*B*; Wilcoxon sign-ranked test: early MC: 3 Hz, mean change in ΔF/F ± SD: = 0.024 ± 0.067, *p* = 1.2×10^−11^; 5 Hz = 0.023 ± 0.078, *p* = 3.0×10^−8^; late MC: 3 Hz, 0.081 ± 0.31, *p* = 1.1×10^−9^; 5 Hz = 0.072 ± 0.4103, *p* = 7.9×10^−3^; early sTC: 3 Hz = 0.17 ± 0.15, *p* = 2. 6×10^−40^; 5 Hz = 0.19 ± 0.25, *p* = 2.6×10^−27^; late sTC: 3 Hz = 0.26 ± 0.32, *p* = 3.4×10^−27^; 5 Hz = 0.13 ± 0.43, *p* = 5.6×10^−9^). However, there was again high variance in the effect of frequency within a given cell, and the net increase in excitation across all cells was smaller in MCs than in sTCs. Together these results indicate that, while increasing inhalation frequency modestly enhances net excitation, repeated odorant sampling reorganizes MC and sTC responses by differentially affecting different cells.

Reformatting of population responses could be a result of changes across all cells, or it could occur across only a subset of cells. To address this we looked at whether reformatting of MC or sTC responses occurred as a result of changes in particular response types. We broke the datasets into groups based on initial response magnitude within the first second of odor presentation. In MCs these groups were strongly responding cells, weakly responding cells, and suppressive cells. In sTCs, owing to the very few suppressive cells seen, we broke the dataset into thirds, consisting of strongly responding, moderately responding, and weakly responding cells. In MCs, cells that initially responded strongly showed a similar response evolution over time across the three inhalation frequencies, consisting of a rapid increase in signal amplitude over the first 2 seconds of odorant presentation, followed by a plateau of responses over the final six seconds (Fig. 4*C*). Among initially suppressive cells, a similar initial increase in amplitude (i.e., greater suppression) followed by a plateau (or even a gradual decrease/slight lessening of suppression in 1 and 3 Hz conditions) was seen. The high variance, especially among cells at 1 Hz, points to substantial evolution of signal amplitude for cells that were initially suppressive. Finally, weakly excited MCs showed a clear frequency-dependence in the evolution of their responses over time, with responses to 5 Hz inhalation increasing four-fold from the first to the last second, compared to approximately three-fold increases at 1 Hz and 3 Hz. Furthermore, at 1 and 3 Hz frequencies, responses showed a gradual increase in amplitude over the 8 s odorant presentation, and even at 5 Hz, where responses plateaued, this plateau was not reached until 4-5 seconds into the odorant presentation. These results suggest that weakly excited MCs may contribute more to population reformatting over time than either strongly excited or suppressed cells, although the high variance in each of the groups, especially among the suppressive cell group, indicates that there is still substantial diversity at the level of individual cells.

In contrast to MCs, sTC response amplitudes showed similar patterns of the percentage change in response over time across all groups (strongly, moderately, and weakly excited cells). In all three groups, responses at 5 Hz rose noticeably over the initial ~two seconds, before gradually declining during the final 5-6 s of the odor presentation, suggestive of adaptation. Responses at 1 and 3 Hz showed a prolonged, but weaker, increase in amplitude over time, before finally falling within the last time bin. The prolonged increase meant that 1 Hz responses showed a higher percentage change than responses across the same cells in the 5 Hz condition (unpaired t-test, t = 3.12, df=8, *p* = 0.014). Thus, MC and sTCs contribute differently to sampling-dependent reformatting of their odor representations, with sTC responses changing in a similar manner over time, regardless of their initial response magnitude.

Glomeruli are the functional unit of odor representations at the level of OB inputs, and each MC has a single apical dendrite that branches heavily in one glomerulus and is the MC’s sole source of excitatory synaptic input (Mombaerts et al., 1996; Schoppa and Westbrook, 2001; Wachowiak et al., 2004; Wachowiak and Shipley, 2006; Economo et al., 2016). We have previously shown that temporally diverse response patterns, including suppressive and excitatory responses, are prevalent in glomerular MC signals (Economo et al., 2016). However, it is possible that different MCs arising from the same glomerulus (i.e., ‘sister’ MCs; (Dhawale et al., 2010)) may show distinct effects of repeated sampling or inhalation frequency, which could ultimately obscure the decorrelation among sister cells. Thus, to further investigate how inhalation frequency impacts odor representations, we imaged MC calcium signals from glomeruli of the dorsal OB.

We found that, as with somatic imaging, glomerular MC responses (n = 15-41 per field of view) were more diverse at higher inhalation frequencies compared to low (Fig. 5*A, B*). Suppressive responses were prevalent, occurring in 472/1164 (40.5%) of all glomerulus-odorant pairs at 1 Hz inhalation, with slightly lower percentages at high inhalation frequencies: 451/1164 (38.7%) at 3 Hz, and 391/1164 (33.6%) at 5 Hz. These percentages are similar to our MC soma data presented above, as well as to those reported earlier using a 2 Hz inhalation frequency (Economo, Hansen, & Wachowiak, 2016). We also observed frequency-dependent changes in responsiveness as well as response polarity: from 1 to 3 Hz, 109/692 (15.7%) glomerulus-odor pairs switched from excited to suppressed, and from 1 to 5 Hz, 111/692 (16.0%) switched from excited to suppressed. Responses that switched from suppressed to excited were more common by a factor of ~2: from 1 to 3 Hz, 114/472 (24.2%) switched from suppressed to excited, and from 1 to 5 Hz, 176/472 (37.3%) switched from suppressed to excited. These percentages are roughly similar to the percentages of MC somas that switched polarity with increasing inhalation frequency.

Other aspects of the glomerular signals differed notably from the MC soma signals. For example, glomerular signals showed less decorrelation at the end of the 8 s odorant presentation compared to somas (Fig 5*C*; t-test, *p* = 0.0037), and they also showed a statistically significant effect of inhalation frequency on the amount of decorrelation, with responses at 1 Hz (*r =* 0.90) greater than those at 3 Hz (*r* = 0.75, Mann-Whitney U-test: *p* = 0.022, n = 11 mice) and 5 Hz (*r* = 0.78; Mann-Whitney U-test: *p* = 0.036, n = 11 mice). Also, glomerular signals imaged with GCaMP6s appeared to show a greater difference in decorrelation between low and high inhalation frequencies compared to the GCaMP6f data (Fig. 5*C*). With both reporters, higher inhalation frequencies resulted in greater decorrelation, though only the result for GCaMP6s was significant (Kruskal-Wallis test: 6s, *χ^2^* = 6.862,12, *p* = 0.032; 6f, *χ^2^* = 2.682,15, *p* = 0.26). Overall, these data point to a similar amount of decorrelation in MC dendrites as in their somas.

Finally, to examine how the evolution of MC and sTC responses might contribute to odor identity coding during repeated sampling, we performed principal components analysis on the somatic population data and measured the Euclidean distance between population response patterns to different odorants as a function of sampling time at each frequency. At 1 Hz inhalation, MC and sTC populations paralleled each other in terms of the dynamics and magnitude of the change in Euclidean distance between different odor representations (Fig. 6*A*).

However, at 3 and 5 Hz inhalation, MC populations diverge from sTC populations midway through the 8 s odor presentation, after which the distance between MC odor representations remains high, while that for sTC representations actually begins to decrease (Fig. 6*B, C*). These results indicate that the population representation of odorants by MCs becomes more distinct with repeated sampling at higher inhalation frequencies compared to low.

Figure 6D demonstrates this effect from an example MC dataset by displaying the change over time of odor responses in principal component space across the three inhalation frequencies. Odor representations grow further apart in the same 8 second block at higher inhalation frequencies compared to low (Fig. 6*D*). In contrast, the same plot in an example sTC imaging experiment (Fig. 6*E*) shows a quick evolution of the signal in PC space (top), followed by periodic “swings” in trajectories corresponding to responses to individual inhalations. This result is qualitatively similar to that seen in ‘synthetic’ populations compiled from multiple MTC recordings (Bathellier et al., 2008). Thus, much of the increase in Euclidean distance between odor representations stems from the initial burst of signal early in the odor presentation, which is commonly seen among sTCs but not MCs (Fig. 2*C, D*). Altogether, the results from this analysis support a role for high-frequency inhalation in generating more distinct odor representations over the course of repeated sampling among MCs, with a smaller impact on odor representations among sTCs.

## Discussion

Previous studies have demonstrated that a single inhalation of odorant is sufficient to robustly encode odorant identity, and that rodents can use information sampled in a single sniff to direct odor-guided behaviors (Rinberg et al., 2006; Kepecs et al., 2007; Wesson et al., 2008b; Wesson et al., 2008a). However, animals routinely sample odorants with bouts of high-frequency sniffing that can last multiple seconds (Vanderwolf, 1992; Thesen et al., 1993; Verhagen et al., 2007; Wesson et al., 2008a). Despite its being a fundamental aspect of active odor sampling, little is known about how such repeated sampling facilitates odor-guided behaviors or impacts odor representations. Earlier studies in diverse species have found that odor representations evolve with sustained odorant stimulation over a period of seconds (Friedrich and Laurent, 2001; Verhagen et al., 2007). Here, by imaging from many cells simultaneously and by targeting defined projection neuron subtypes, we have gained new insights into this process in the mammalian olfactory system. We found that repeated odorant sampling impacts odor representations by MCs and CCK+ sTCs differently, with MCs showing a reformatting of the initial odor representation that increased with increasing inhalation frequency, while the sTC odor representation remained relatively consistent across repeated samples and across frequencies. These results further support the idea that MCs and TCs constitute distinct functional pathways for early odor information processing, and suggest that the reformatting of MC odor representations by high-frequency sniffing may serve to enhance the discrimination of similar odors.

Our results are largely consistent with predictions from a previous study that relied on whole-cell recordings from presumptive MT cells (Díaz-Quesada et al., 2018), which found that individual MT cells showed diverse and frequency-dependent changes in excitation with repeated sampling at 1, 3 and 5 Hz. Likewise, they are consistent with recordings of presumptive MT cells in awake mice, which found substantial changes in MT cell spiking patterns over the first few inhalations of odorant (Carey and Wachowiak, 2011; Patterson et al., 2013; Díaz-

Quesada et al., 2018). The fact that we observed cell type-specific and frequency-dependent reformatting of MT cell odor representations that was sustained over a longer time-period of up to 8 seconds in anesthetized mice suggests that the changes seen in awake mice likely result from ‘bottom-up’ effects related to changes in the dynamics of incoming sensory input, as opposed to modulation by centrifugal inputs related to behavioral state.

Imaging from a relatively large population of cells simultaneously allowed us to gain insights into the nature of the reformatting of the MC overall population response. We observed that initial response magnitude influenced how MCs responded over time, with initially weakly-responding cells exhibited the most variability (and greatest differentiation) of responses over the course of odorant presentation. Weakly-responding MCs showed the greatest change in activity over the 8 s odor presentation and, at high frequency, these cells substantially increased their response magnitudes compared to lower frequencies (Fig. 4C). This result is consistent with findings from slice preparations that MT cell spike output increases with increasing input frequency over a similar range (Balu et al., 2004). This finding contrasts with that for CCK+ sTCs, where time- and frequency-dependent changes in excitation were more systematic and occurred more uniformly across the population. The differences between these two cell types may point to their differing abilities to encode parameters of odorant stimuli.

Our finding that MC responses evolved in the face of changing inhalation frequency substantially more than sTCs supports the idea that these two subpopulations subserve distinct roles in odor information coding. In the context of active odor sampling, sTCs may convey a reliable, sniff-by-sniff snapshot of an animal’s odor environment, while MCs may have the capacity to encode multiple aspects of odor information and do so flexibly, as a function of the animal’s sampling behavior. MC activity patterns may also reflect distinct stimulus features at different times during a sniff bout, similar to what occurs in MCs of the zebrafish OB (Friedrich et al., 2004; Friedrich and Laurent, 2004), and relatedly, as occurs with repeated tastant sampling among neurons of gustatory cortex (Katz et al., 2001).

The circuit mechanisms underlying the frequency-dependent reformatting of MC odor representations remain to be determined. Inhibition from granule cells onto MC lateral dendrites has been proposed to decorrelate ‘sister’ MCs on a faster time-scale (Dhawale et al., 2010), and this process may also work across repeated sampling. However, the fact that we observed similar, albeit somewhat smaller, time- and frequency-dependent changes in patterns of MC activity imaged across OB glomeruli, each of which contains the apical dendrites of multiple ‘sister’ MCs, points to circuit mechanisms in the glomerular layer as playing an important role in this reformatting. One such mechanism may be interglomerular inhibitory circuits that are differentially engaged depending on the overall pattern of glomerular excitation relative to a given MC (Economo et al., 2016; Díaz-Quesada et al., 2018). sTCs are thought to be less subject to this inhibition (Christie et al., 2001; Christie and Westbrook, 2006; Fukunaga et al., 2012; Phillips et al., 2012; Jordan et al., 2018b), consistent with our findings that these cells show more uniform responses over time and across inhalation frequency.

An important caveat to the comparison of sTCs and MCs in this study is that we targeted sTCs defined by their expression of CCK, which constitutes a subset of the entire TC population (Seroogy et al., 1985; Liu and Shipley, 1994; Tobin et al., 2010). We have previously shown that CCK+ sTCs show shorter-latency and simpler excitatory responses to odorants even relative to the larger tufted cell population ((Economo et al., 2016; Short and Wachowiak, 2019)). This difference may explain why, in our previous study that relied simply on soma depth to define MT subtype, we did not observe differences in prevalence of suppressive responses or impact of inhalation frequency on excitability of tufted and mitral cells (Díaz-Quesada et al., 2018). However, other studies have observed marked differences in tufted and mitral cell response dynamics using soma depth as an identifying criterion (Fukunaga et al., 2012). Future experiments using additional genetic or anatomical criteria – for example, cortical projection target or expression of other transmitter receptors – will be important in better understanding the nature of diversity in the response properties of OB output neurons.

In awake, behaving animals, inhalation frequencies routinely reach 10 Hz or even higher, which is substantially faster than the highest frequency used here. However, even the 5 Hz inhalation frequencies used in this study—which approach technical limits of our artificial inhalation approach—show dramatic impacts on odor representations when contrasted with lower frequencies (Carey and Wachowiak, 2011; Díaz-Quesada et al., 2018). Additionally, while brief bouts of high frequency inhalations may reach 10-12 Hz, sustained bouts (lasting 2 or more seconds) rarely stay so high (Wesson et al., 2008a). At these higher frequencies we might expect even greater decorrelation of initial response patterns, leading to rapid population-level reformatting of odorant responses. We also predict that such bouts could lead to higher proportions of suppressive responses in MCs, based on the modest increase in the prevalence of suppression seen when inhalation frequencies increased from 1 to 5 Hz. Such a result could approach levels of suppression seen in awake mice (e.g., 46% in (Kollo et al., 2014)).

The diverse changes in MT cell excitability contrast with those of odorant responses among OSNs, where high-frequency inhalation more uniformly attenuates OSN inputs due to adaptation (Verhagen et al., 2007). How this adaptation leads to a more complex reformatting of MTC response patterns, including both decreases and increases in excitation, is unclear. Recently, by imaging glutamate signals onto MTC dendrites, we observed greater than expected diversity in the dynamics and magnitude of excitatory input onto MTCs during repeated sampling an anesthetized and awake mice (Moran et al., 2019). Thus, patterns of OSN-driven excitation onto MT cells may be more diverse than previously thought. In future experiments it will be important to image simultaneously from OSNs and MTCs to gain further insight into the mechanisms underlying this reorganization of odor representations as a function of sampling behavior.

## Acknowledgments

We thank S. Burton, A. Moran, S. Short, and I. Youngstrom for helpful comments, G. Vasquez for help with virus injections, and J. Ball and R. Kummer for help with animal husbandry and histological processing. This work was supported by funding from NIH (F32DC015389 to TPE and DC006441 to MW).

## References

Abraham NM, Spors H, Carleton A, Margrie TW, Kuner T, Schaefer AT (2004) Maintaining accuracy at the expense of speed: stimulus similarity defines odor discrimination time in mice. Neuron 44:865–876.

Abraham NM, Egger V, Shimshek DR, Renden R, Fukunaga I, Sprengel R, Seeburg PH, Klugmann M, Margrie TW, Schaefer AT, Kuner T (2010) Synaptic Inhibition in the Olfactory Bulb Accelerates Odor Discrimination in Mice. Neuron 65:399–411.

Balu R, Larimer P, Strowbridge BW (2004) Phasic stimuli evoke precisely timed spikes in intermittently discharging mitral cells. J Neurophysiol 92:743–753.

Bathellier B, Buhl DL, Accolla R, Carleton A (2008) Dynamic Ensemble Odor Coding in the Mammalian Olfactory Bulb: Sensory Information at Different Timescales. Neuron 57:586–598.

Carey RM, Wachowiak M (2011) Effect of sniffing on the temporal structure of mitral/tufted cell output from the olfactory bulb. J Neurosci 31:10615–10626.

Christie J, Schoppa N, Westbrook G (2001) Tufted cell dendrodendritic inhibition in the olfactory bulb is dependent on NMDA receptor activity. J Neurophysiol 85:169–173.

Christie JM, Westbrook GL (2006) Lateral Excitation within the Olfactory Bulb. J Neurosci 26:2269–2277.

Courtiol E, Hegoburu C, Litaudon P, Garcia S, Fourcaud-Trocme N, Buonviso N (2011) Individual and synergistic effects of sniffing frequency and flow rate on olfactory bulb activity. J Neurophysiol 106:2813–2824.

Dhawale AK, Hagiwara A, Bhalla US, Murthy VN, Albeanu DF (2010) Non-redundant odor coding by sister mitral cells revealed by light addressable glomeruli in the mouse. Nat Neurosci 13:1404–1412.

Díaz-Quesada M, Youngstrom IA, Tsuno Y, Hansen KR, Economo MN, Wachowiak M (2018) Inhalation frequency controls reformatting of mitral/tufted cell odor representations in the olfactory bulb. Journal of Neuroscience 38:2189–2206.

Economo MN, Hansen KR, Wachowiak M (2016) Control of mitral/tufted cell output by selective inhibition among olfactory bulb glomeruli. Neuron 91:397–411.

Eiting TP, Wachowiak M (2018) Artificial inhalation protocol in adult mice. Bio-protocol 8.

Friedrich R, Habermann C, Laurent G (2004) Multiplexing using synchrony in the zebrafish olfactory bulb. Nat Neurosci 7:862–871.

Friedrich RW, Laurent G (2001) Dynamic optimization of odor representations by slow temporal patterning of mitral cell activity. Science 291:889–894.

Friedrich RW, Laurent G (2004) Dynamics of olfactory bulb input and output activity during odor stimulation in zebrafish. J Neurophysiol 91:2658–2669.

Fukunaga I, Berning M, Kollo M, Schmaltz A, Schaefer Andreas T (2012) Two Distinct Channels of Olfactory Bulb Output. Neuron 75:320–329.

Gong S, Zheng C, Doughty ML, Losos K, Didkovsky N, Schambra UB, Nowak NJ, Joyner A, Leblanc G, Hatten ME, Heintz N (2003) A gene expression atlas of the central nervous system based on bacterial artificial chromosomes. Nature 425:917–925.

Griff ER, Mafhouz M, Chaput MA (2008) Comparison of Identified Mitral and Tufted Cells in Freely Breathing Rats: II. Odor-Evoked Responses. Chem Senses 33:793–802.

Haddad R, Lanjuin A, Madisen L, Zeng H, Murthy VN, Uchida N (2013) Olfactory cortical neurons read out a relative time code in the olfactory bulb. Nat Neurosci 16:949–957.

Jordan R, Kollo M, Schaefer AT (2018a) Sniffing fast: paradoxical effects on odor concentration discrimination at the levels of olfactory bulb output and behavior. eNeuro 5:e0148–18.

Jordan R, Fukunaga I, Kollo M, Schaefer AT (2018b) Active sampling state dynamically enhances olfactory bulb odor representation. Neuron 98:1214–1228.

Katz DB, Simon SA, Nicolelis MA (2001) Dynamic and multimodal responses of gustatory cortical neurons in awake rats. J Neurosci 21:4478–4489.

Kepecs A, Uchida N, Mainen ZF (2007) Rapid and precise control of sniffing during olfactory discrimination in rats. J Neurophysiol 98:205–213.

Kollo M, Schmaltz A, Abdelhamid M, Fukunaga I, Schaefer AT (2014) ‘Silent’mitral cells dominate odor responses in the olfactory bulb of awake mice. Nat Neurosci 17:1313.

Laska M (1990) Olfactory sensitivity to food odor components in the short-tailed fruit bat, Carollia perspicillata (Phyllostomatidae, Chiroptera). J Comp Physiol A 166:395–399.

Liu WL, Shipley MT (1994) Intrabulbar associational system in the rat olfactory bulb comprises cholecystokinin-containing tufted cells that synapse onto the dendrites of GABAergic granule cells. J Comp Neurol 346:541–558.

Madisen L, Garner AR, Shimaoka D, Chuong AS, Klapoetke NC, Li L, Van Der Bourg A, Niino Y, Egolf L, Monetti C (2015) Transgenic mice for intersectional targeting of neural sensors and effectors with high specificity and performance. Neuron 85:942–958.

Mombaerts P, Wang F, Dulac C, Chao SK, Nemes A, Mendelsohn M, Edmondson J, Axel R (1996) Visualizing an olfactory sensory map. Cell 87:675–686.

Moran AK, Eiting TP, Wachowiak M (2019) Diverse dynamics of glutamatergic input underlie heterogeneous response patterns of olfactory bulb mitral and tufted cells *in vivo*. bioRxiv:692574.

Nagayama S, Takahashi YK, Yoshihara Y, Mori K (2004) Mitral and Tufted Cells Differ in the Decoding Manner of Odor Maps in the Rat Olfactory Bulb. J Neurophysiol 91:2532–2540.

Najac M, De Saint Jan D, Reguero L, Grandes P, Charpak S (2011) Monosynaptic and Polysynaptic Feed-Forward Inputs to Mitral Cells from Olfactory Sensory Neurons. J Neurosci 31:8722–8729.

Patterson MA, Lagier S, Carleton A (2013) Odor representations in the olfactory bulb evolve after the first breath and persist as an odor afterimage. PNAS 110:E3340–3349.

Phillips ME, Sachdev RNS, Willhite DC, Shepherd GM (2012) Respiration Drives Network Activity and Modulates Synaptic and Circuit Processing of Lateral Inhibition in the Olfactory Bulb. J Neurosci 32:85–98.

Rinberg D, Koulakov A, Gelperin A (2006) Speed-accuracy tradeoff in olfaction. Neuron 51:351–358.

Rothermel M, Brunert D, Zabawa C, Díaz-Quesada M, Wachowiak M (2013) Transgene Expression in Target-Defined Neuron Populations Mediated by Retrograde Infection with Adeno-Associated Viral Vectors. J Neurosci 33:15195–15206.

Rygg AD, Van Valkenburgh B, Craven BA (2017) The influence of sniffing on airflow and odorant deposition in the canine nasal cavity. Chem Senses 42:683–698.

Schoppa NE, Westbrook GL (2001) Glomerulus-specific synchronization of mitral cells in the olfactory bulb. Neuron 31:639–651.

Seroogy KB, Brecha N, Gall C (1985) Distribution of cholecystokinin-like immunoreactivity in the rat main olfactory bulb. J Comp Neurol 239:373–383.

Short SM, Wachowiak M (2019) Temporal dynamics of inhalation-linked activity across defined subpopulations of mouse olfactory bulb neurons imaged in vivo. eNeuro 6:0189–0119.

Taniguchi H, He M, Wu P, Kim S, Paik R, Sugino K, Kvitsani D, Fu Y, Lu J, Lin Y, Miyoshi G, Shima Y, Fishell G, Nelson Sacha B, Huang ZJ (2011) A Resource of Cre Driver Lines for Genetic Targeting of GABAergic Neurons in Cerebral Cortex. Neuron 71:995–1013.

Thesen A, Steen JB, Doving KB (1993) Behaviour of dogs during olfactory tracking. J Exp Biol 180:247–251.

Tobin VA, Hashimoto H, Wacker DW, Takayanagi Y, Langnaese K, Caquineau C, Noack J, Landgraf R, Onaka T, Leng G, Meddle SL, Engelmann M, Ludwig M (2010) An intrinsic vasopressin system in the olfactory bulb is involved in social recognition. Nature 464:413–417.

Uchida N, Mainen ZF (2003) Speed and accuracy of olfactory discrimination in the rat. Nat Neurosci 6:1224–1229.

Uchida N, Kepecs A, Mainen ZF (2006) Seeing at a glance, smelling in a whiff: rapid forms of perceptual decision making. Nat Rev Neurosci 7:485–491.

Vanderwolf CH (1992) Hippocampal activity, olfaction, and sniffing: an olfactory input to the dentate gyrus. Brain Research 593:197–208.

Verhagen JV, Wesson DW, Netoff TI, White JA, Wachowiak M (2007) Sniffing controls an adaptive filter of sensory input to the olfactory bulb. Nat Neurosci 10:631–639.

Wachowiak M (2011) All in a sniff: olfaction as a model for active sensing. Neuron 71:962–973.

Wachowiak M, Cohen LB (2001) Representation of odorants by receptor neuron input to the mouse olfactory bulb. Neuron 32:723–735.

Wachowiak M, Shipley MT (2006) Coding and synaptic processing of sensory information in the glomerular layer of the olfactory bulb. Semin Cell Dev Biol 17:411–423.

Wachowiak M, Denk W, Friedrich RW (2004) Functional organization of sensory input to the olfactory bulb glomerulus analyzed by two-photon calcium imaging. PNAS 101:9097–9102.

Wachowiak M, Economo MN, Díaz-Quesada M, Brunert D, Wesson DW, White JA, Rothermel M (2013) Optical Dissection of Odor Information Processing In Vivo Using GCaMPs Expressed in Specified Cell Types of the Olfactory Bulb. J Neurosci 33:5285–5300.

Welker WI (1964) Analysis of sniffing in the albino rat. Behavior 22:223–244.

Wesson DW, Donahou TN, Johnson MO, Wachowiak M (2008a) Sniffing behavior of mice during performance in odor-guided tasks. Chem Senses 33:581–596.

Wesson DW, Carey RM, Verhagen JV, Wachowiak M (2008b) Rapid Encoding and Perception of Novel Odors in the Rat. PLoS Biology 6:e82.

Youngentob SL, Mozell MM, Sheehe PR, Hornung DE (1987) A quantitative analysis of sniffing strategies in rats performing odor detection tasks. Physiol Behav 41:59–69.

Zhao K, Dalton P, Yang GC, Scherer PW (2006) Numerical modeling of turbulent and laminar airflow and odorant transport during sniffing in the human and rat nose. Chem Senses 31:107–118.

